# KMAP: Kmer Manifold Approximation and Projection for visualizing DNA sequences

**DOI:** 10.1101/2024.04.12.589197

**Authors:** Chengbo Fu, Einari A. Niskanen, Gong-Hong Wei, Zhirong Yang, Marta Sanvicente-García, Marc Güell, Lu Cheng

## Abstract

Identifying and illustrating patterns in DNA sequences is a crucial task in various biological data analyses. In this task, patterns are often represented by sets of kmers, the fundamental building blocks of DNA sequences. To visually unveil these patterns, we could project each kmer onto a point in two-dimensional (2D) space. However, this projection poses challenges due to the high-dimensional nature of kmers and their unique mathematical properties. Here, we established a mathematical system to address the peculiarities of the kmer manifold. Leveraging this kmer manifold theory, we developed a statistical method named KMAP for detecting kmer patterns and visualizing them in 2D space. We applied KMAP to three distinct datasets to showcase its utility. KMAP achieved a comparable performance to the classical method MEME, with approximately 90% similarity in motif discovery from HT-SELEX data. In the analysis of H3K27ac ChIP-seq data from Ewing Sarcoma (EWS), we found that BACH1, OTX2 and ERG1 might affect EWS prognosis by binding to promoter and enhancer regions across the genome. We also found that FLI1 bound to the enhancer regions after ETV6 degradation, which showed the competitive binding between ETV6 and FLI1. Moreover, KMAP identified four prevalent patterns in gene editing data of the AAVS1 locus, aligning with findings reported in the literature. These applications underscore that KMAP could be a valuable tool across various biological contexts. KMAP is freely available at: https://github.com/chengl7-lab/kmap.

## INTRODUCTION

DNA sequence serve as the primary carrier of genetic information. In diverse research contexts, investigators aim to uncover patterns within DNA sequences, a pursuit central to various applications. Among the most prominent of these is the study of transcription factor (TF) DNA-binding specificity. Researchers employ methods like SELEX-seq (Jolma et al., 2013; Kinzler & Vogelstein, 1990; Tuerk & Gold, 1990), ChIP-seq (Johnson, Mortazavi, Myers, & Wold, 2007; Wei et al., 2010) to determine the binding specificities of TFs. SELEX-seq, an *in vitro* technique, typically examines a single TF at a time, whereas H3K27ac ChIP-seq that targets active regulatory regions, an *in vivo* approach, allows for the investigation of the binding of multiple TFs. The DNA sequences obtained from these methodologies provide crucial biological insights into TF activities, such as their DNA-binding specificities. Furthermore, DNA sequencing is also utilized to study the effect of gene editing protocols, where researchers are interested in discovering the editing patterns. Therefore, the DNA sequence encapsulate a wide array of biological information, varying according to the specific application in focus.

To extract the biological information from DNA sequences, researchers usually convert these sequences into kmers. The kmer distribution mirrors the underlying biological information, allowing for the identification of potential patterns. For instance, in SELEX-seq data, kmers relevant to the DNA-binding specificity of a certain TF may cluster together. Similarly, in gene editing datasets, multiple clusters might represent editing patterns. Therefore, it would be very convenient for researchers to explore the kmer distribution by projecting each kmer to a point in the two-dimensional (2D) Euclidean space. However, this task presents two main challenges. Firstly, the kmer space is extremely large. The total number of possible kmers is 4^*k*^, which grows exponentially as *k* increases. It brings a huge computation load to project so many points to the 2D space when *k* is large. Furthermore, human eyes struggle to discern real patterns among a dense collection of data points in 2D space, where meaningful signals may be overwhelmed by noisy points. Secondly, the discrete nature of kmers has introduced special topological properties to the kmer space, which forms the kmer manifold. There are intrinsic conflicts between the kmer manifold and the Euclidean space, whereas current dimensionality reduction methods are designed for the continuous Euclidean space. Classical motif discovery algorithms such as MEME (Bailey & Elkan, 1994), HOMER (Heinz et al., 2010), STREME (Bailey, 2021) address the issue of kmer explosion by filtering kmers that are relevant to a motif pattern. These selected kmers are used to construct a position weight matrix representing the motif. However, this approach typically focuses on individual motifs one at a time, failing to capture the full spectrum of patterns in the kmer distribution. For example, the relative strengths between different patterns are not directly observable. Displaying all patterns within a single figure would be significantly more informative, offering a comprehensive view of the underlying biological information.

Due to the special properties of the kmer manifold, researchers rarely get satisfactory results in kmer visualization. Kruppa et al. (2017) proposed visualizing the kmer distribution as bubbles anchored on a pyramid, but this method lacked intuitiveness. Yi et al. (2021) randomly scattered kmers in a 2D space, yet this approach failed to reveal underlying motif patterns. Classical dimensionality reduction methods designed for Euclidean space such as Principal Component Analysis (PCA; Hotelling, 1933) and Multidimensional Scaling (MDS; Torgerson, 1952) are linear dimensionality reduction methods, which cannot cope with peculiarities of the kmer manifold. Other methods such as tSNE (Van der Maaten & Hinton, 2008) and UMAP (McInnes, Healy, & Melville, 2018) could handle data from a nonlinear perspective, but they face the issue of gradient explosion, especially when the space is discrete and there are a large number of duplicate points in the input. Yuan et al. (2019) alleviate this problem by learning a high dimensional continuous embedding (*d*=300) using a supervised neural network that matches a bag of kmers with a single TF label, which learns the embeddings of TFs instead of kmers. Their approach, however, is not suitable for the kmer manifold visualization task here, which is inherently an unsupervised task. Deterministic unsupervised neural networks like autoencoder (Kramer, 1992) suffer from the identical mapping problem that identical kmers would be projected to the same point, which leads to the loss of the density information. Probabilistic unsupervised neural networks such as variational autoencoders (VAE; Khemakhem, Kingma, Monti, & Hyvarinen, 2020), offer a promising alternative. VAE has been used for motif discovery from ATAC-seq data (Kshirsagar, Yuan, Ferres, & Leslie, 2022), though their potential for kmer manifold visualization remains untapped.

In this study, we aim to project each kmer onto a point in 2D space, to provide an intuitive visualization of the kmer distribution. The key to achieving this objective is to explore the unique properties of the kmer manifold. As far as we are aware, there is not a proper theoretical formalization of the kmer manifold in the field of Bioinformatics. To fill this gap, we have built up the mathematical theories for describing the kmer manifold. Leveraging this theoretical framework, we examined the probability distribution of kmers, introduced the concept of Hamming ball, and developed a motif discovery algorithm, such that we could sample relevant kmers to depict the full kmer manifold. After that, we performed transformations to the kmer distances based on the kmer manifold theory to mitigate the inherent discrepancies between the kmer manifold and the 2D Euclidean space. Finally, we developed the KMAP visualization algorithm to project kmers to 2D space by extending the UMAP framework.

To demonstrate its utility, we applied KMAP to various datasets, including SELEX-seq, ChIP-seq and gene editing data. The comparison with the established MEME algorithm (Bailey & Elkan, 1994) on a comprehensive benchmark SELEX-seq dataset shows that KMAP generates similar motifs as MEME. Specifically, in the analysis of Ewing Sarcoma (EWS) data, we not only reproduced the original findings (Lu et al., 2023) about GGAA-repeats in the motif of ETS Variant Transcription Factor 6 (ETV6), but also unveiled new insights. We discovered ETV6 could also bind to AGGG repeats and likely inhibited the binding of BACH1, OTX2, ERG1 to the promoter and enhancer regions. Furthermore, in our analysis of the gene editing data of the AAVS1 locus, KMAP identified four distinct editing patterns. Remarkably, these patterns correspond to the top 4 patterns reported in the original publication (Sanvicente-García et al., 2023), reinforcing KMAP’s capability to discern and visualize complex genetic patterns.

## RESULTS

### Kmer manifold theory and KMAP workflow

We have built up the mathematical theories for studying the kmer manifold, where the most important conclusions are illustrated in Fig. 1A-C. Let us denote a kmer by 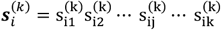. Here 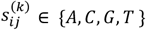 and *k* is the length of the kmer. There are 4^*k*^ unique kmers in the kmer space. By selecting a kmer as the origin of the kmer space, we could partition all kmers into *k* + 1 orbits (Fig. 1A), where kmers of the *i*-th orbit (*i* = 0, 1, 2, …, *k*) have *i* mutations compared with the origin. In other words, the Hamming distance between each kmer in the *i*-th orbit and the origin is *i*. The kmer manifold Ω^(*k*)^ is jointly defined by the origin, the metric (Hamming distance) and all kmers in the kmer space. In Supplementary Note 1, we derived the count of kmers in each orbit of Ω^(*k*)^:

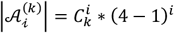

We have proved that this function is an unimode function of the orbit index *i* (Fig. 1B), with the mode located near 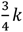. As illustrated by Fig. 1A, orbits 5, 6, 7 have the densest points, where 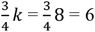. We define kmers of orbits within a radius of *r*^(*k*)^ as a **Hamming Ball** (red circle in Fig. 1A) to represent kmers that are similar to the origin. As shown in Fig. 1B, the count of kmers within the Hamming ball is relatively small (∼0.4%) for *k*=8 and *r*^(*k*)^=2. Fig. 1C provides the empirical distribution of the ratio between the empirical probability and the theoretical probability for all Hamming balls from a random DNA sequence, which we term as **Hamming ball ratio**. The empirical probability of a Hamming ball is given by the proportion of kmers within the Hamming ball out of all kmers in the input DNA sequence, while the theoretical probability is given by (see Sec.5 of Supplementary Note 1):

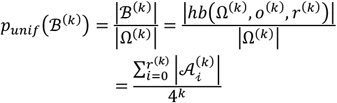

We fit a Gaussian distribution to the Hamming ball ratios over all kmers with the mean fixed to 1 and use it as the null distribution. Based on practical experiences, we set the p-value threshold to 1 × 10 ^−10^ for a Hamming ball ratio to be considered as significant. Detailed description of the kmer manifold theory is given in Supplementary Note 1, where we also discuss reverse complements of kmers and how to treat them in practice.

The motif discovery algorithm (Fig. 1D) is based on the kmer manifold theory, particularly, the null distribution of the Hamming ball ratio. We have proved that the kmer manifold is isotropic, i.e. we could generate the whole kmer space by centering on any kmer (as origin) in the manifold. Therefore, a motif can be represented by a Hamming ball, where the origin is termed as the **consensus sequence** of the motif. From real sequencing data, we can count kmers and calculate the actual probability of a Hamming ball centered on a top kmer, which is tested against the null distribution. Hamming balls with a p-value less than the significance threshold are kept as motifs. It is difficult to visualize the kmer space as it is extremely large, e.g. only 0.4% kmers land in the Hamming ball and majority kmers are random kmers in Fig. 1A. Here we term kmers within a motif Hamming ball as **motif kmers** and other kmers as **random kmers**. We perform sampling of kmers for visualization after motif discovery. Half of the sampled kmers are motif kmers, while the other half are random kmers. kmers from different motifs are pooled and sampled with the weights given by their counts. Random kmers are sampled in a similar manner to reflect the background noise. Detailed description of the motif discovery algorithm is provided in Supplementary Note 2.

**Figure 1:**
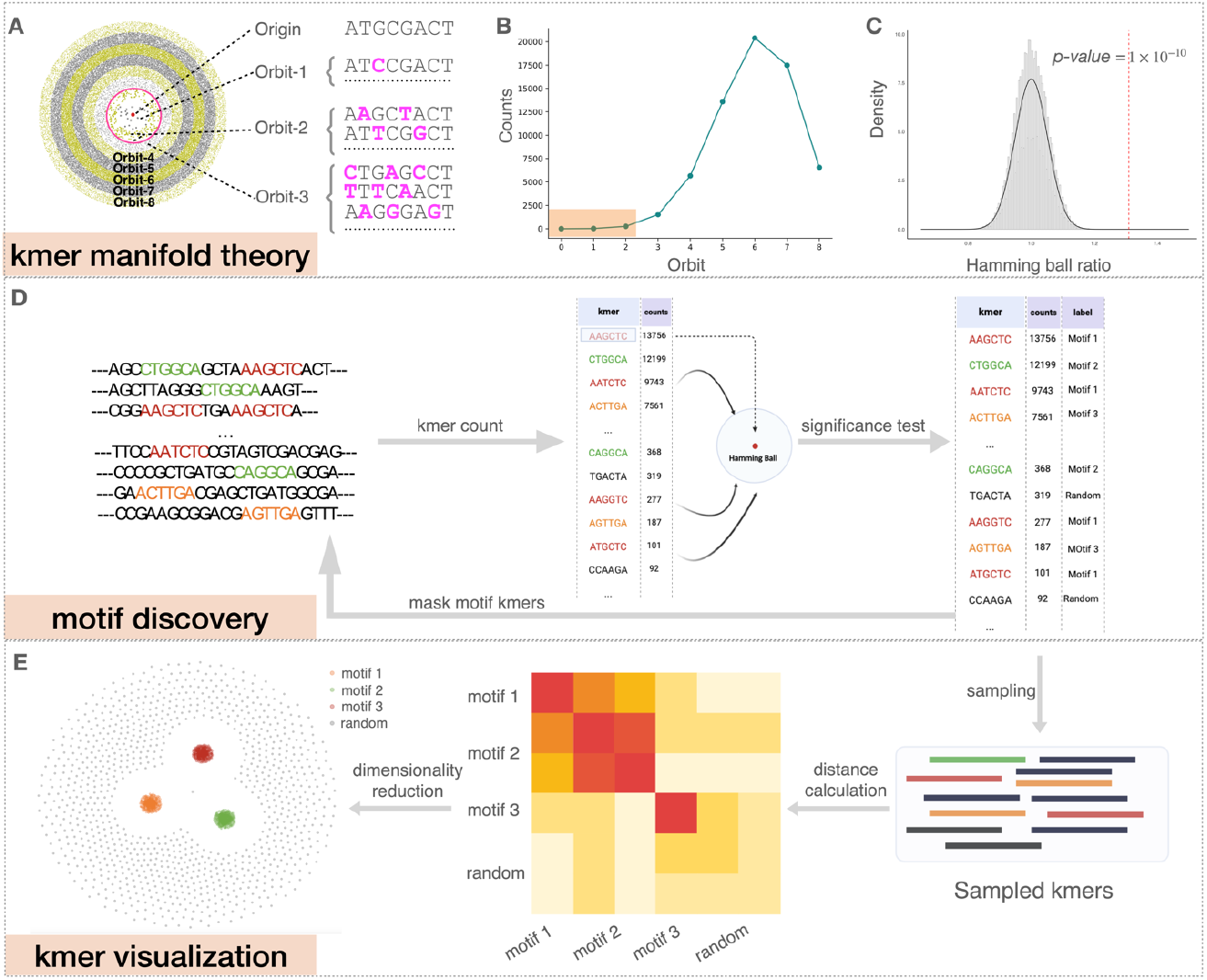
KMAP workflow. (A) Schematic illustration of the kmer manifold for *k*=8. Each point represents a unique kmer. Orbit-*i* consists of kmers with *i* mutations to the origin, i.e. the Hamming distance from the kmer to the origin is *i*. kmers in the *i-*th orbit are uniformly scattered in the *i-*th ring, where each ring has an equal width. kmers within the red circle forms the Hamming ball centered on the origin with a radius *r*^(*k*)^ = 2. (B) kmer counts of each orbit. The rectangle highlights the kmer counts of orbits in the Hamming ball. (C) Null distribution of Hamming ball ratio. The histogram is generated by taking all Hamming ball ratios from a random DNA sequence of 100000 bp. The experiment is repeated 10 times and a Gaussian distribution is fitted to the obtained ratios with the mean fixed to 1. The fitted Gaussian distribution is used as the null distribution, where the vertical dashed line indicates the significant ratio corresponding to a p-value of 1 × 10^−10^. (D) The motif discovery workflow. We first count the kmers, then test the Hamming ball centered on the top kmer, after that we mask all motif kmers from the input DNA sequence and repeat the process iteratively until no motif can be found. (E) kmer visualization algorithm. 2500 motif kmers and 2500 random kmers are sampled for the visualization. The Hamming distance matrix of the sampled kmers is smoothed and further utilized for dimensionality reduction.

We visualize the sampled kmers *X* = (***x***_1_, ***x***_2_, ⋯, ***x***_*N*_) by projecting them to 2D space (Fig. 1E). First, we calculate the Hamming distance matrix between the kmers. Then we compute the smoothed distance matrix by pulling neighboring kmers and repulsing distant kmers. Neighboring kmers are pulled closer by the following distance transformation:

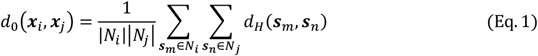

where ***s***_*m*_ and ***s***_*n*_ are one of the 20 nearest neighbors of ***x***_*i*_ and ***x***_*j*_, respectively. Here *N*_*i*_ and *N*_*j*_ denote the 20 nearest neighbors of ***x***_*i*_ and ***x***_*j*_. *d*_*H*_(***s***_*m*_, ***s***_*n*_) denotes the Hamming distance between ***s***_*m*_ and ***s***_*n*_.

Distant kmers are repulsed away by the following transformation:

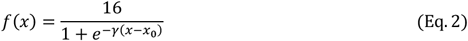

where *x* is the transformed distance (Eq. 1), *γ* = 0.2*k* − 0.2 controls the curvature of the transformation, 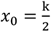 is the change point parameter and *k* is the length of the input kmer. *x*_0_ is the rough boundary between Hamming ball and the outer orbits. According to Remark 1.1 in Supplementary Note 1, the expected distance between two random kmers is 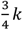, i.e. a kmer in the Hamming ball and a kmer in the outer orbits. Hence, we choose 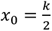 as the rough boundary.

These transformations try to mitigate the intrinsic conflicts between Hamming distance and Euclidean distance. As shown in Fig. 2A, all six kmers have exactly 1 mutation from the consensus sequence, so we arrange them on a circle of radius 1. However, the Hamming distance between any pair of kmers is 2. There is no way to arrange the kmers on the circle that satisfies these constraints. The most intuitive solution is to place them as a hexagon on the circle. Although the Euclidean distances between diagonal kmers (black edges) are 2, the Euclidean distances between adjacent kmers (green edges) and semi-diagonal kmers (blue edges) are 1 and 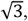 which are less than the designed Hamming distances 2. As a result, directly utilizing the Hamming distances leads to inferior results. Fig. 2A (right panel) shows the effect of distance smoothing, which pulls neighboring kmers and repulses distant kmers, as well as introduces randomness to the distances. We demonstrate the Hamming distance matrix and its transformation on a toy example with 3 motifs in Fig. 2B. KMAP visualizations based on the original and transformed Hamming distance matrices are provided in Fig. 2C. It can be seen that the transformation pulls motif kmers closer and repulses random kmers further, which improves the visualization effect.

**Figure 2:**
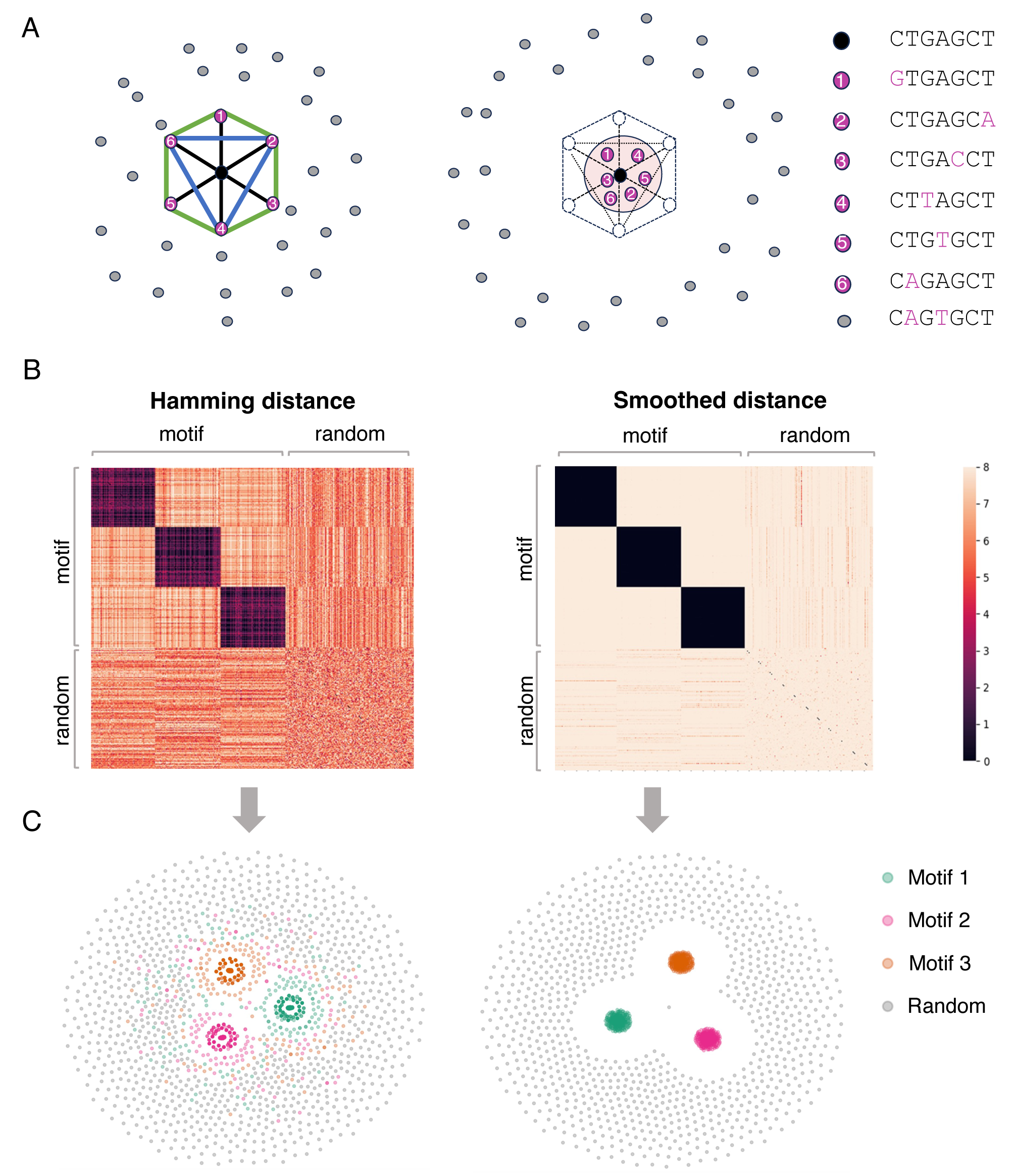
Peculiarities of kmer manifold. (A) Six motif kmers (orange dots) with one mutation from the origin (black dot) placed as a hexagon on a circle of radius 1, while random kmers (gray dots) are placed outside. The Hamming distance between each pair of motif kmers is 2. The Euclidean distance between the dots are 2, 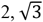, 1 for diagonal (black line), semi-diagonal (blue line) and adjacent kmers (green line). The right panel shows the schematic effects of Hamming distance transformations (Eq. 1 & 2), where motif kmers are pulled closer and random kmers are repulsed further. (B) Toy example. The left panel shows the Hamming distance matrix of a kmer (*k*=8) dataset with 3 motifs, as highlighted by the black blocks. Within a motif, the Hamming distance ranges from 0 to 4. The right panel shows the transformed Hamming distance matrix. After transformation, the distances between the motif kmers are reduced, while the distances between motif and random kmers become larger. (C) KMAP visualizations based on the original Hamming distance matrix (left) and the transformed Hamming distance matrix (right).

The KMAP visualization algorithm projects the high dimensional kmers to 2D space, given the transformed Hamming distance matrix. We denote the *i*-th kmer as **x**_*i*_ and its 2D embedding as ***w***_*i*_ = (*w*_*i*0_ *w*_*i*1_),where *i* = 1, 2, …, *N*. We use the same cross entropy loss function as UMAP for dimensionality reduction:

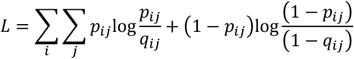

where *p*_*ij*_ is the similarity probability of the high dimensional data, given by:

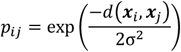

and *q*_*ij*_ is the similarity probability of the low dimensional embeddings, given by:

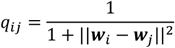

It is obvious that the loss function reaches its minimum when *p*_*ij*_ = *q*_*ij*_ for all *i* ≠ *j*. By optimizing the loss function, we try to find embeddings that generate *q*_*ij*_ as close as *p*_*ij*_, such that the low dimensional embeddings could represent the high dimensional manifold. We use the gradient descent algorithm for the optimization. The gradient is given by:

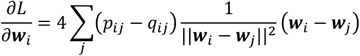

We notice that the term ||***w***_*i*_ – ***w***_*j*_||^2^ can easily go to zero in the optimization, which causes gradient exploding. We add the following diffusion terms to *w*_*j*_ to avoid this problem.

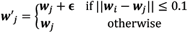

where **ϵ** = (ϵ_0,_ ϵ_1_) and ϵ_0,_ ϵ_1_ ∼ *N*(0, 0.01^2^) are two independent Gaussian samples.

## HT-SELEX data analysis

We demonstrate KMAP’s performance in motif discovery on a public high-throughput SELEX (HT-SELEX) dataset (Jolma et al., 2013), which contains 461 TFs. We analyze 1273 samples of SELEX round 3-6 for these TFs, which have stronger motif signals compared with round 1-2. Fig. 3A shows motif logos for 4 example TFs: BHLHB2, MAFK, MEF2D and NFKB2 given by KMAP and classical methods such as MEME, STREME, DREME. It can be seen that motifs given by different methods are similar. We further compared the results of KMAP and MEME, where only the top motif is compared. For each sample, we calculated the precision and recall scores from the consensus sequences given by KMAP and MEME using the following formulas:

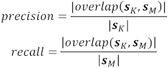

where | ⋅ | takes the length of a given sequence, ***s***_*K*_ and ***s***_*M*_ denote the consensus sequences provided by KMAP and MEME, respectively. Fig. 3B shows the distribution of the precision and recall scores over all samples. The precision and recall medians between KMAP and MEME are 82% and 92%, which suggests a high consistency between KMAP and MEME. Note that the precision is generally smaller than the recall, which suggests that KMAP generates longer motifs than MEME.

**Figure 3:**
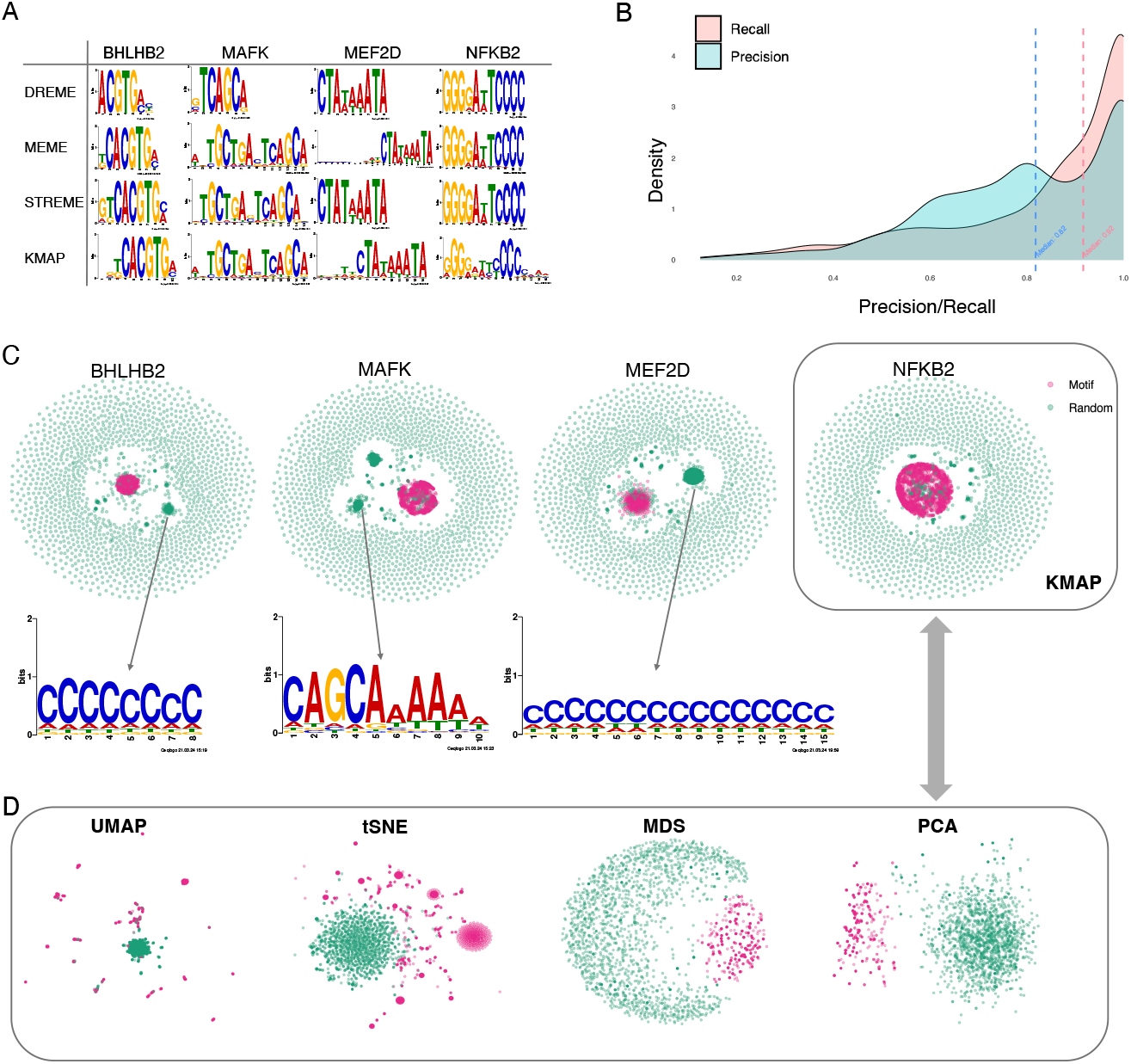
Benchmark on HT-SELEX data. (A) Motif logos of four example TFs given by DREME, MEME, STREME, KMAP. (B) Distribution of precision and recall scores between MEME and KMAP. The scores are calculated for 1273 samples (SELEX round 3-6), with the corresponding medians highlighted by the dashed vertical lines. (C) KMAP visualizations of example TFs. Motif and random kmers are highlighted in red and green, respectively. The logos illustrate exemplary secondary motifs of TFs. (D) kmer visualizations of NFKB2 based on UMAP, tSNE, MDS, PCA. KMAP visualization of NFKB2 is highlighted in the rectangle above.

We show the KMAP 2D embeddings of kmers for the example TFs in Fig. 3C, where secondary motifs could be observed for BHLHB2, MAFK, MEF2D. These secondary motifs are distinct to the major motifs. Fig. 3D illustrates the dimensionality reduction results of UMAP, tSNE, MDS, PCA, which are generated using the same set of kmers of NFKB2. It can be seen that motif kmers (red dots) do not form a clear cluster in the PCA and MDS embeddings. Due to gradient exploding, tSNE and UMAP stop after a few iterations. We can see that motif kmers are scattered around and random kmers form a cluster, which is likely due to initialization. KMAP provides the most intuitive representation of the kmer manifold (Fig. 3C, rightmost panel), where Hamming balls form clusters in the center and random kmers are placed in the peripheral space.

### Ewing Sarcoma data analysis

We analyzed ChIP-seq data from an Ewing Sarcoma (EWS) study (Lu et al., 2023). Lu et al. have found that ETV6 promotes the development of EWS by competitively binding to the binding sites of FLI1, which is also confirmed in another study (Gao et al., 2023). Lu et al. prepared ETV6-dTAG A673 or EW8 cells, from which ETV6 could be rapidly degraded by adding a small molecule called dTAG^V^-1, whereas ETV6 was intact if DMSO was added. From ChIP-seq data targeting ETV6 or FLI1 in the parental A673 cells (WT), KMAP identified GGAA-repeats as the strongest motif for FLI1 and ETV6 (Fig. 4A). From ChIP-seq data of A673 ETV6-dTAG cells with dTAG^V^-1 treated, where ETV6 was degraded, KMAP found the GGAA-repeats motif disappeared in ChIP-seq data targeting ETV6 (Fig. 4A) but remained in data targeting FLI1. Fig. 4B shows the motif logos generated from the Hamming ball of GGAA-repeats (16bp) and AGGG-repeats (13bp) motifs. These findings have confirmed the original conclusion that ETV6 and FLI1 competitively bind to GGAA-repeats on the genome. It is worth mentioning that ETV6 also binds to AGGG-repeats, which is the second strongest motif in WT. The “GGAAGG” motif with a single “GGAA” repeat can be observed in the ETV6-KO sample, which suggests the existence of a small amount of ETV6 in the ETV6-KO sample. Since ETV6 is both an oncogenic gene and a repressor, we are interested in its target genes, which might inhibit the development of EWS.

**Figure 4:**
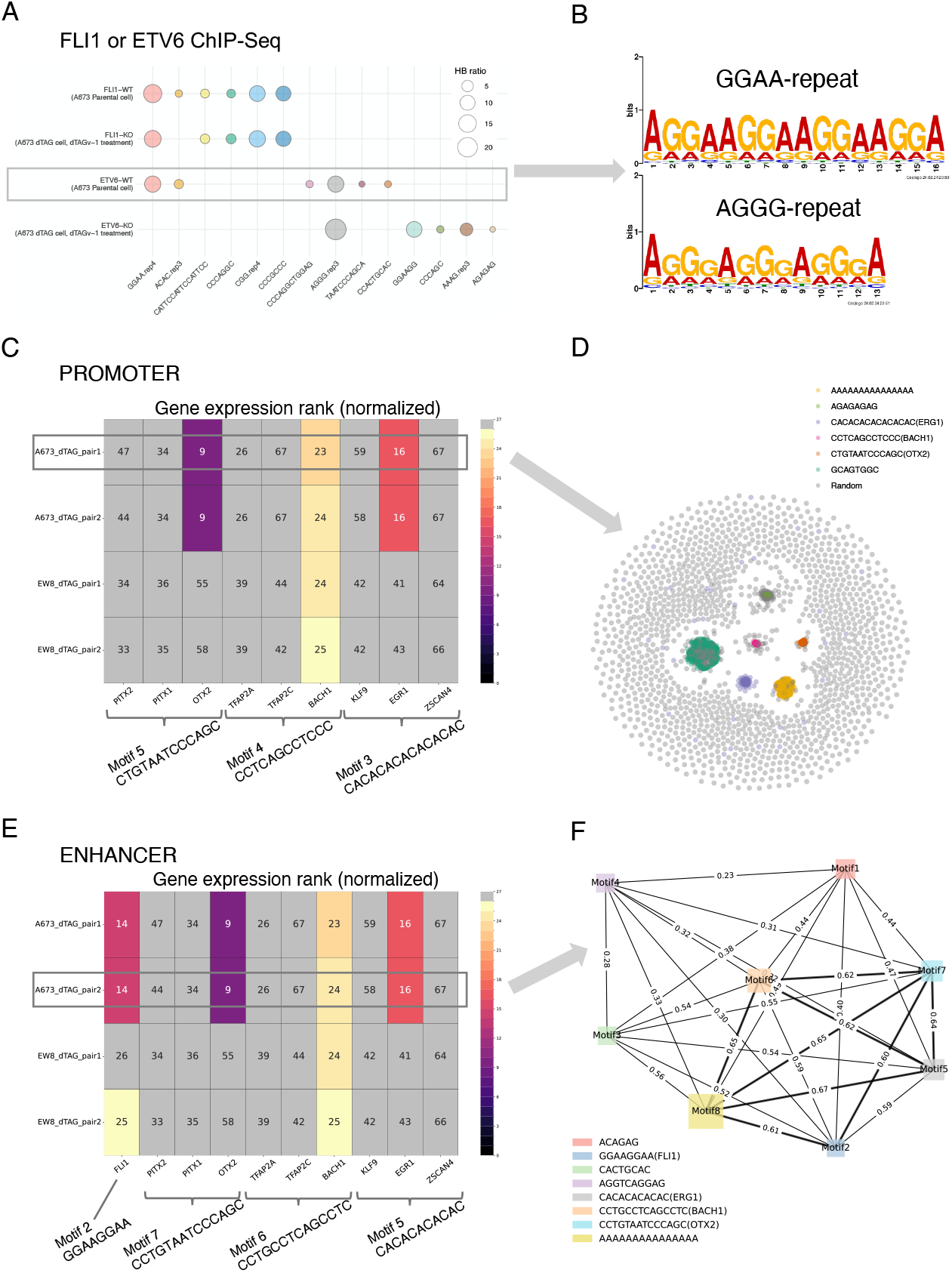
Ewing Sarcoma data analysis. (A) Motifs identified by KMAP from ChIP-seq data targeting FLI1 or ETV6. WT refers to A673 parental cells. KO refers to dTAG^V^-1 treated ETV6-dTAG cells derived from A673 parental cells. In the motifs, “rep4” means repeat larger or equal to 4 times and similar rule applies to “rep3”. The circle size indicates the Hamming ball ratio. (B). Motif logos of GGAA and AGGG repeats generated from motif kmers, based on ChIP-Seq peaks of the ETV6-WT sample. (C) Expression levels of potential promoter region associated TFs. Each row is a sample pair, i.e. dTAG cells treated with dTAG^V^-1 (ETV6 degraded) or DMSO (ETV6 intact), where newly gained promoter regions (ETV6 degraded vs intact) are used as the input for KMAP. The columns are the potential TFs of the identified motifs. Note that FIMO returns multiple TFs for a single input consensus sequence. For each pair, gene expressions in the corresponding ETV6 degraded sample (dTAG^V^-1 treated) are ranked and the ranks are normalized to 0-100, where top 25% genes are colored, and the rest genes are gray. The number in each cell shows the normalized rank, where smaller values mean higher expressions. The motif consensus sequences are obtained from the “A673_dTAG_pair1” pair highlighted by the rectangle. (D) KMAP visualization of kmers in the promoter region, based on newly gained ChIP-seq peaks (ETV6 degraded vs intact) from the “A673_dTAG_pair1” pair. (E) Expression levels of potential TFs in enhancer regions. The motif consensus sequences are extracted from the “A673_dTAG_pair2” pair. (D) Co-occurrence network of identified motifs from the “A673_dTAG_pair2” pair. The edge weight indicates the proportion of ChIP-seq peaks that contain both TFs out of all peaks that contain at least one TF. The node size indicates the Hamming ball ratio.

Lu et al. (2023) provided the H3K27ac ChIP-seq data of A673 and EW8 ETV6-dTAG cells, which marked the promoter and enhancer regions of ETV6 target genes. There are 8 samples divided into 4 pairs, each of which contains a control sample with intact ETV6 (DMSO treated) and a ETV6 degraded sample (dTAG^V^-1 treated). Each pair corresponds to a specific combination of the cell line (A673 or EW8) and the replicate, e.g. “A673-dTAG-pair1” refers to A673 dTAG cells with dTAG^V^-1 (ETV6 degraded, replicate 1) or DMSO (ETV6 intact, replicate 1). For each pair, we extracted the H3K27ac ChIP-seq peaks that were gained upon ETV6 degradation. This was done by subtracting peaks of the ETV6 degraded sample from that of the ETV6 intact sample, i.e. dTAG^V^-1 vs DMSO. The newly gained peaks were further classified into promoter and enhancer regions. KMAP identified 8 unique motifs from the promoter regions. We next fed the consensus sequences of the 8 unique motifs to FIMO (Grant, Bailey, & Noble, 2011) to retrieve the corresponding TFs, where one consensus sequence might match several TFs. In total, FIMO identified 3 motifs with available annotations in at least one of the pairs. Assuming the potential TFs were highly expressed, we extracted the gene expressions of the matched TFs in the ETV6 degraded samples (dTAG^V^-1 treated). Here we ranked expressions of all genes and normalized the ranks to 0-100, where smaller numbers indicated higher expressions. Fig. 4C shows the potential TFs binding to the promoter region, where BACH1are identified as a motif in almost all pairs and have a high expression. OTX2 and ERG1 are likely only associated with the A673 dTAG cells. Fig. 4D shows the kmer manifold of the newly gained ChIP-seq peaks (ETV6 degraded vs ETV6 intact) from the A673-dTAG-pair1 pair, where 6 motifs are observed.

Similar analysis of the enhancer regions (Fig. 4E) has identified 8 motifs, 4 out of which have matched TFs after motif identification using FIMO. It can be seen that FLI1 and BACH1 have a high expression in almost all pairs, while OTX2 and ERG1 are only associated with the A673 cells. Fig. 3F illustrates the TF co-occurrence network generated from newly gained ChIP-seq peaks (ETV6 degraded vs intact) from the “A673_dTAG_pair2” pair. This result shows that motif 2, 5, 6, 7, 8 (FLI1, ERG1, BACH1, OTX2, A-repeats) have a high frequency of co-occurrence, while motif 1, 3, 4 occurs less frequently with other motifs.

These findings suggest that (1) ETV6 prevents the binding of BACH1, OTX2 and ERG1 to the promoter and enhancer regions, which may have implications for EWS prognosis (2) ETV6 and FLI1 competitively bind to the enhancer regions more than the promoter regions.

### Gene editing data analysis

We use KMAP to detect editing patterns from gene editing data of the Adeno-Associated Virus Site 1 (AAVS1) locus. Sanvicente-García et al. (2023) used a previously described (Doench et al., 2016; Mali et al., 2013) guide RNA with a 20nt protospacer that targeted the genomic location (chr19:55,115,752-55,115,771; GRCh38/hg38) on the human genome. The Streptococcus pyogenes Cas9 (SpCas9) protein will make a double stranded break (DSB) at three nucleotides upstream of the PAM, after that the broken DNAs are ligated by non-homologous end joining (NHEJ), which leads to different DNA editing results (mainly random small indels). Another cellular repair mechanism involved in the repair of DSB is microhomology-mediated end-joining (MMEJ), resulting in longer deletions led by homologous patterns in both sides of the cleavage. It is interesting to explore if there exist any patterns in the gene editing results, since patterns that lead to a certain repair resolution, allow a higher rate of success to achieve certain outcomes, like efficient knockouts with out-of-frame deletions. From the gene editing results (fastq files) of the aforementioned experiment, we first removed DNA reads that were identical to the reference sequence, which represented unedited DNA. Next, we generated a multiple sequence alignment from the remaining reads using MUSCLE (Edgar, 2004). After that, we calculated the pairwise distance matrix between the aligned reads and performed unsupervised sampling in KMAP (Algorithm 2 of Supplementary Note 2). Given the Hamming distance matrix of the sampled reads (*n*=2696), we directly performed dimensionality reduction using KMAP (Fig. 5A). Based on the 2D embeddings, we identified 6 clusters/patterns using DBSCAN (Ester, Kriegel, Sander, & Xu, 1996). The patterns in our results nicely agree with that of Sanvicente-García et al.’s CRISPR-A platform. Pattern 1 (Fig. 5B) represents sequencing errors due to poly nucleotide and high GC content in the protospacer and its flanking regions. Pattern 5 (Fig. 5C) represent reads from another genomic region near the 3’ end of the reference sequence that is distant to the protospacer. The remaining 4 patterns (Fig. 5D-G) have a one-to-one correspondence with Sanvicente-García et al.’s results, which represent a 12 nucleotides (nt) MMEJ deletion, a 5nt MMEJ deletion, a 3nt CAG insertion adjacent to the cleavage site, 1nt deletion at the cut site, respectively. Out of these 4 patterns, patterns 2 and 3 are the most abundant.

**Figure 5:**
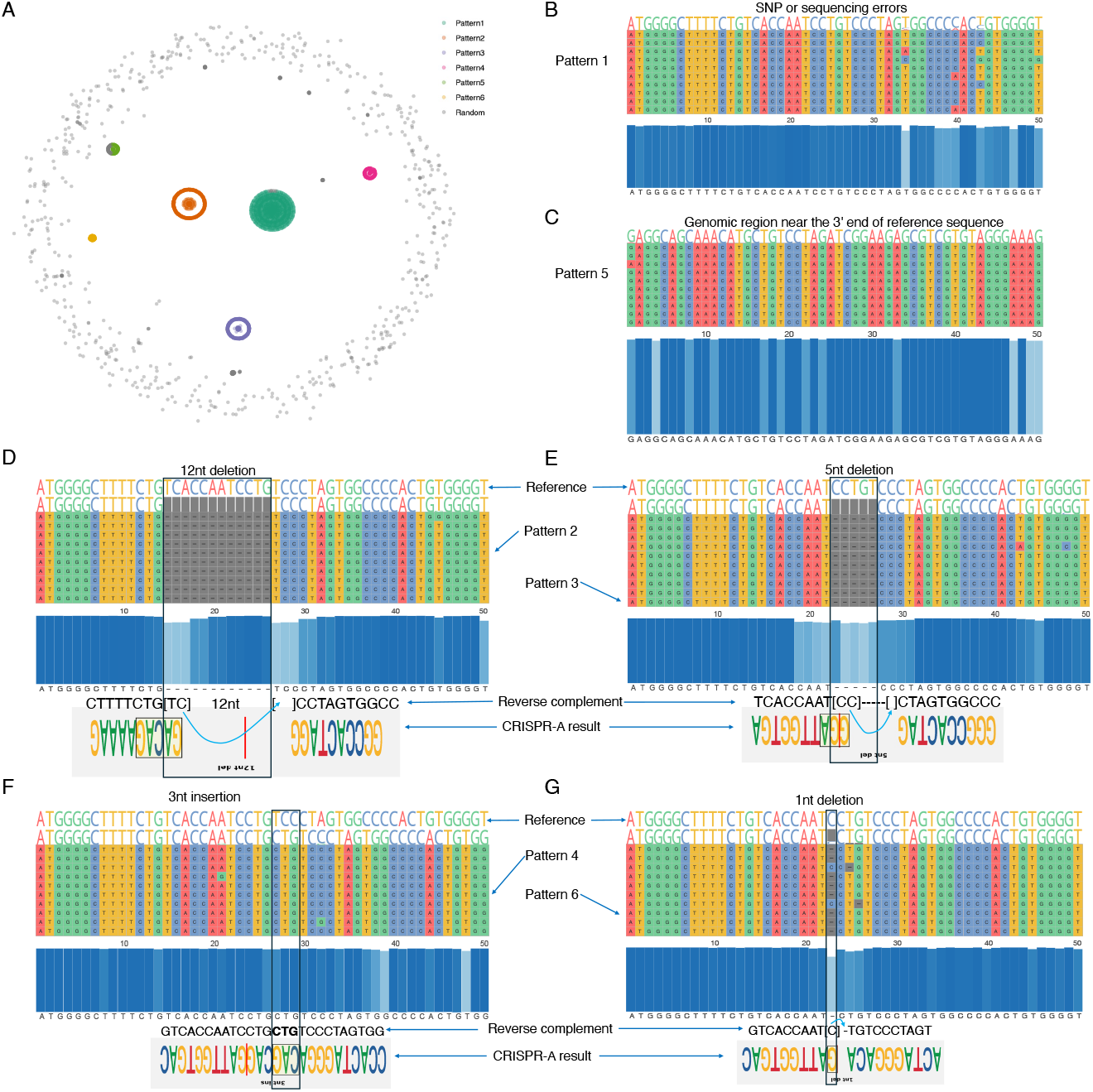
Gene editing patterns. (A) KMAP visualization of aligned DNA sequences. The six clusters are given by DBSCAN based on the 2D embeddings. (B-G) Sequence alignment of each pattern. Each row represents a DNA sequence. Ten sequences from each cluster are shown in the alignment to illustrate the corresponding pattern. The blue panel in the center shows the conservative nucleotide at each position. The top 4 gene editing patterns given by CRISPR-A are provided at the bottom panel for the last 4 patterns (D-G), above which are their reverse complements. Note that although the deletions in our result show 1-3nt shifts compared with CRISPR-A’s result, they are equivalent as the shifted nucleotides can be placed on either side of the deletion in the alignment. Pattern 1 is treated as SNPs or sequencing errors by CRISPR-A. Pattern 5 corresponds to a different genomic region.

## DISCUSSION

kmer is an important object in various DNA sequence studies. A set of kmers, or a Hamming ball in this context, can provide a description of the binding preferences of a given TF. If we turn an input sequence data into a set of kmers, the kmers form clusters that potentially have biological interpretations. An intuitive idea is to project kmers to 2D space to visualize them.

kmers are high-dimensional objects living in the kmer manifold with special properties. We dive deep into the math to build up the theories for the kmer manifold. The kmer manifold is isotropic. By centering on any kmer, the kmer manifold can be partitioned into *k* + 1 orbits, where majority kmers are in orbits with indices close to 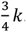. Although the Hamming ball contains a relatively large number of kmers when *k* is large, they only constitute a tiny fraction of the whole kmer manifold. We derived the null distribution of the Hamming ball ratio and used it for motif discovery. The reverse complement operation adds an additional layer of symmetry to the kmer manifold. We spend lots of efforts discussing how a Hamming ball is partitioned into four parts due to the reverse complement operation in Supplementary Note 1, which greatly helps the real data treatment.

There exist intrinsic conflicts between the kmer manifold and the Euclidean space. The metric of the kmer manifold is the Hamming distance, which possesses counter-intuitive properties as shown in Fig. 2A. We performed distance smoothing to harmonize the conflicts between Hamming distance and Euclidean distance. The idea is that the neighbors of a motif kmer are likely still neighbors of another motif kmer, but their distances to a random kmer are still near 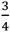*k*, which allows us to further scale the distance to repulse random kmers. The discreteness of the Hamming distance often induces zero to the denominator of a term in the gradient, which causes the gradient exploding problem and pre-termination of the optimization. Adding a small diffusion term to identical embeddings can effectively solve this problem.

Beyond the distance smoothing and diffusion term, there is another difference between KMAP and UMAP. In UMAP, a point specific scaling parameter is used to pull distant points such that the scaled distances to other points follow the standard Gaussian distribution. This treatment has a side effect in the kmer manifold, where distant kmers (random kmers) are pulled closer and form a cluster. In KMAP, we do not perform point specific scaling, which preserves the global structure of the kmer manifold. Therefore, random kmers are scattered in the peripheral space. Due to the huge volume (4^*k*^) of the kmer manifold, we had to sample a subset of kmers for the visualization. The bottleneck is the calculation of the distance matrix, which contains *n*^2^ elements for *n* input kmers.

The kmer manifold theory has enabled us to study the kmer distribution in different biological contexts. We have shown that KMAP provided similar results as MEME in the HT-SELEX data. In the Ewing Sarcoma data, we found that BACH1, OTX2 and ERG1 -motif containing promoters and enhancers are masked by ETV6. This suggest that BACH1, OTX2 and ERG1 could be involved in alleviating EWS progression, which could be tested by activating the expression of BACH1, OTX2 and ERG1. Activating of BACH1, OTX2 and ERG1-motif containing promoters and enhancers could also result from increase in EWS-FLI activity at the enhancer regions upon ETV6 degradation. Indel distribution visualization is very important for the gene editing community. Selection of gRNAs that induce low complexity distributions which are absent of in-frame indels are highly desirable to maximize knock out implementation. In the gene editing data, KMAP has identified the major editing patterns. We believe KMAP could be used for other tasks such as analyzing RNA-protein interactions (CLIP-seq), ATAC-seq etc.

There are open questions from the method perspective. We use a heuristic to determine the motif length *k*. If a sequence and its subsequences are identified as motif sequences for three consecutive lengths *k* − 1, *k, k* + 1, we use *k* as the final motif length. However, it is unclear how a Hamming ball changes when the motif length changes from *k* to *k* + 1. New theories are needed to depict the transformation of the kmer manifold from *k* to *k* + 1. Given the success of such a theory, we could investigate the possibilities of projecting the *k* and *k* + 1 kmer manifolds to 2D and 3D jointly, such that we could visualize the transformation process. This will help to address the motif length determination problem. Another direction worth further investigation is how to characterize the kmer manifolds for two different conditions, e.g. WT and KO, and how to compare the kmer manifolds and get the differentially expressed Hamming balls.

In summary, we have developed a computational framework KMAP for visualizing kmers in different biological contexts.

## METHOD

### KMAP analysis workflow

The following workflow was used both for HT-SELEX and EWS data analysis. For KMAP motif discovery, we set the following default parameters: *k* = 5, 6, …, 16, the ratio threshold of significance was 1× 10^−10^, the reverse complement mode was on. Results of different kmer lengths were merged to generate the final motifs (Hamming ball), each of which could have a different length. KMAP motif discovery also provided the co-occurrence matrix of the motifs. For visualization, 5000 data points were sampled (supervised sampling). For KMAP dimensionality reduction, we used 20 neighbors for distance smoothing, set the learning rate to 0.01 and used 2500 iterations for the optimization. The outcome of KMAP dimensionality reduction was the 2D embeddings of input kmers.

kmers in the motif (Hamming ball) were used for position weight matrix (PWM) calculation and logo generation, by “sites2meme” and “ceqlogo” commands in MEME Suite (version= 5.0.5) (Bailey, Johnson, Grant, & Noble, 2015), as well as LogoMaker (version=0.8) (Tareen & Kinney, 2020)

### HT-SELEX data analysis

The data was downloaded from European Nucleotide Archive under the accession number ERP001824. We selected 1273 samples of SELEX cycle 3 to 6 for the benchmark comparison of KMAP and MEME. We limited the motif length to range from 5 to 16 and only picked the top motif in the comparison both for KMAP and MEME. Single nucleotide repetitive motifs such as ‘AAAAA’ or ‘CCCCC’ were removed from KMAP output. Exact match is used to identify the overlap between KMAP and MEME consensus sequences.

The following samples are used for the illustration of Figure 1A: BHLHB2 (Cycle 4, 137919 sequences), MAFK (Cycle 6, 662800 sequences), MEF2D (Cycle 4, 580243 sequences) and NFKB2 (Cycle 5, 132941 sequences). Default parameters were used in DREME (Bailey, 2011), STREME (Bailey, 2021) analyses. The motif length was set to range from 5 to 16 and the strongest motif is selected for illustration.

The scikit-learn package (Pedregosa et al., 2011) were used for PCA, MDS, tSNE, UMAP analysis, where default parameters were used. kmers were converted to one-hot vectors to be used as the input of PCA. The Hamming distance matrix of input kmers was used as the input for MDS, tSNE, UMAP.

### EWS data analysis

All data were downloaded from project GSE181554 in NCBI’s Gene Expression Omnibus (GEO). Detailed list of files and the meta information, as well as their names shown in the figures, are provided in the Supplementary Table 1.

For ChIP-seq/cut-tag analysis, SRA files were converted to fastq files using the “fastq-dump” command in SRAtools (version=3.0.5). Reads were aligned to the human genome (hg19) using Bowtie2 (version=2.5.1) (Quinlan & Hall, 2010) using the ‘—very sensitive’ preset collection of parameters. Samtools (version=1.6) (Danecek et al., 2021) was used to convert bam files into sam files. ChIP-seq peaks were called using MACS2 (version=2.2.7.1) (Zhang et al., 2008) with default parameters by comparing with the corresponding input DNA control. The derived peaks (bed files) were converted to a fasta file using bedtools (version=2.31.0) (Quinlan & Hall, 2010).

The differential peaks (H3K27ac) were derived by subtracting peaks of the ETV6 intact sample from that of ETV6 degraded sample, using the “subtract” command of bedtools. Promoter regions were identified as (-2000bp, +200bp) of transcription start sites, whereas other regions were treated as enhancer regions. Peaks were classified into these two categories using the “annotatePeak” function from the ChIPseeker R package (Yu, Wang, & He, 2015), which used the annotation file of hg19 human genome from “TxDb.Hsapiens.UCSC.hg19.knownGene” package (Carlson & Maintainer, 2015). The corresponding DNA sequences on the genome of these peaks were fed to KMAP analysis. We deleted motifs that were full of “A” s in KMAP results, which were presumed to be noise.

Given the consensus sequences of KMAP motifs, FIMO in MEME Suite were used to identify the TFs of the motifs from JASPAR database (Castro-Mondragon et al., 2022), with default p-value threshold of 1e-4. The expression levels of all genes (TPM column from the quantified gene expression file in the original publication, accession: GSM5505965) were ranked for each sample to generate the quantiles (ranks were scaled to 0-100%) for all relevant TFs. For each sample, a gene is treated as a high-expressing gene if its quantile is less than 25%.

To derive the co-occurrence matrix of different motifs, we scan for all motif consensus sequences on each input read, then record the co-occurrence of all identified motif pairs. We get the co-occurrence count matrix after scanning all reads. For each pair of motifs, we normalized its co-occurrence count by dividing the number of reads that contain either motif. The normalized co-occurrence matrix is fed to NetworkX (version=2.2) (Hagberg, Swart, & S Chult, 2008) to generate the co-occurrence network.

### Gene editing data analysis

Fastq file, Protospacer sequence, reference sequence were obtained from CRISPR-A website (https://synbio.upf.edu/crispr-a/documentation.html#fourth, access date 7.2.2024).

#### Reference sequence

GCTCCAGGAAATGGGGGTGTGTCACCAGATAAGGAATCTGCCTAACAGGAGGTGGGGGTTAGACCCAATATCAGGAGACTAGGAAGGAGGAGGCCTAAGGATGGGGCTTTTCTGTCACCAATCC TGTCCCTAGTGGCCCCACTGTGGGGTGGAGGGGACAGATAAAAGTACCCAGAACCAGAGCCACATTAACCGGCCCTGGGAATATAAGGTGGTCCCAGCTCGGGGACACAGGATCCCTGGAGGCAGCAAACATGCTGTCCT.

#### Protospacer

GGGGCCACTAGGGACAGGAT

Filtered reads were aligned using MULCLE (v5) (Edgar, 2004) with the “super5” command. Multiple sequence alignment was visualized using ggmsa (version=1.4.0) package (Zhou et al., 2022). DBSCAN (Ester et al., 1996) in sklearn (version=1.3.0) (Pedregosa et al., 2011) was used to identify clusters from 2D embeddings, with parameters *eps=3* and *min_sample=200*.

Since there are only ∼20000 reads and the read length are too long to be treated as a kmer (*k*=250), we performed unsupervised sampling (Algorithm 2, Supplementary Note 2) on the data. We calculated the pairwise Hamming distance matrix between the aligned reads, from which we derived the smoothed distance matrix by taking the average of 600 nearest neighbors. We then fitted a Gaussian distribution to smoothed distances, which was given by *N*(46.5, 14.9^2^). We kept distances that were less than *µ* − 2σ=15.56, which presumably correspond to motif kmers/sequences. Sequences involved in the filtered distances were the output of the unsupervised sampling. Hamming distance matrix of the sampled DNA sequences was directly used in the dimensionality reduction.

## DATA ACCESS

The KMAP package is freely available on https://github.com/chengl7-lab/kmap. Analysis scripts are deposited on https://github.com/Dionysos-o/kmap/tree/main/kmap_paper.

## ACKNOWLEDGMENTS

We would like to acknowledge Prof. Aki Vehtari from Aalto University for his comments on the method development and manuscript writing. Additionally, we extend our gratitude to M.Sc. Fanduo Li from Meituan (Shanghai, China), for his assistance with the kmer manifold theory in the supplementary materials. We acknowledge the computational resources provided by the Aalto Science-IT project.

## CONTRIBUTION

CF and LC developed the mathematical theory and implemented the KMAP software. CF conducted the analyses of all experiments. EN and GW provided advice about EWS data analysis. MG and MS provided advice about gene editing data analysis. ZY provided advice about the visualization algorithm development. All authors contributed to the manuscript writing. LC conceptualized and supervised the whole project.

